# Mutation-induced reshaping of protein conformational dynamics revealed by a coarse-grained modeling framework

**DOI:** 10.64898/2026.03.29.715126

**Authors:** Byung Ho Lee, Domenico Scaramozzino, Serena Piticchio, Laura Orellana

**Affiliations:** Department of Oncology-Pathology, Karolinska Institutet, SE-171 77 Stockholm, Sweden

## Abstract

Disease-related missense mutations reshape protein conformational energy landscapes, thereby altering biological function. However, mechanistically linking sequence variation to changes in conformational dynamics remains challenging for both experimental and computational approaches. Here, we introduce an internal-coordinate-based, essential-dynamics-refined elastic network model (ICed-ENM) that improves the physical fidelity of normal modes while capturing subtle mutation-induced side-chain effects and preserving computational efficiency. By constraining bond-length and bond-angle fluctuations and refining mode subspaces against experimentally observed collective motions, ICed-ENM provides a stable, structure-encoded description of intrinsic protein dynamics. Building on this framework, we developed a systematic mutation-scanning analysis that quantifies mutation impact as changes in vibrational entropy, providing a dynamic measure of mutation-induced redistribution within conformational energy landscapes. Validation against all-atom molecular dynamics simulations demonstrates that residues predicted as mutation hot spots induce substantial reshaping of free-energy landscapes, consistent with altered intrinsic conformational dynamics. Extending this analysis across a curated protein structure dataset reveals global patterns of mutation sensitivity across diverse structural and physicochemical contexts. Notably, these trends align with large-scale public mutation datasets, suggesting that our framework captures features relevant to pathogenic variation. Together, ICed-ENM and the associated mutation-scanning pipeline provide a scalable and mechanistically interpretable strategy to identify mutation-sensitive regions and substitutions, offering deeper insight into how sequence variation reshapes functional conformational landscapes.

## 1. Introduction

Substitution of a single amino acid on a protein sequence, known as a missense mutation, can have a critical impact on its biological function, even leading to various pathologies from mutation-driven cancers (1, 2) to neurodegenerative disorders, including Alzheimer’s and Parkinson’s disease (3). Such mutations may contribute to subtly alter the conformational energy landscape, as well as directly impair active sites or compromise structural stability, collectively affecting their biological functions (4). However, the precise pathway through which these mutations exert their influence can be challenging to determine. In particular, the allosteric effect is difficult to interpret, as it involves mutations that exert distant influences on the active site through long-range dynamic communication (5, 6). Consequently, a deep understanding of the impact of missense mutations on conformational energy landscapes is essential for elucidating the mechanistic foundations of oncogenesis.

Numerous efforts have been made to capture conformational changes and, by extension, to assess the influence of mutations in protein dynamics. Molecular dynamics (MD) simulations are a well-established approach for exploring the conformational energy landscape of proteins (7). Although advances in computational power (8, 9) and conformational sampling strategies (10–13) have greatly improved the ability of MD simulations to capture large-scale transitions and rare conformational states, intrinsic limitations remain. These include a strong dependence on force field accuracy and high computational cost (14), which still hinder the ability to capture transitions across high-energy barriers and to detect subtle shifts in the conformational energy landscape of mutated proteins.

As a faster, more computationally efficient alternative, coarse-grained (CG) modeling has become widely adopted to explore intrinsic protein dynamics. Among these approaches, elastic network models (ENMs), which represent proteins as mechanical networks of mass points (typically Cα atoms) connected by harmonic springs, have been extensively applied to investigate collective, low-frequency motions (15–17) and conformational transitions (18–20). To account for mutation effects, several extensions have incorporated residue-specific interactions into ENMs. For example, a sequence-sensitive ENM has been proposed to refine the elastic network by modulating spring constants based on atomic contact surfaces and amino-acid identities (21). In parallel, machine-learning frameworks have emerged that integrate ENM-derived dynamic descriptors with structural and evolutionary features to estimate mutation-induced perturbations in protein stability or flexibility (22, 23). However, these approaches provide limited means to directly assess mutation-induced reshaping of protein conformational energy landscapes. Furthermore, machine-learning predictors have intrinsic limitations, including their strong dependence on the availability and quality of training datasets and their lack of clear physical interpretability (24, 25).

To address these limitations, we develop a systematic framework for quantifying mutation-induced perturbations of protein conformational energy landscapes. As a central component of this framework, we introduce an internal-coordinate-based, essential-dynamics-refined elastic network model (ICed-ENM) designed to capture reliable protein dynamics and sensitively detect subtle changes in dynamics induced by single-residue substitutions. Unlike conventional ENMs formulated in Cartesian coordinate (CC) space, ICed-ENM operates in internal coordinates (ICs) and incorporates dynamic information derived from all-atom MD simulations, thereby embedding atomistic flexibility and anharmonic features into a coarse-grained framework. Leveraging ICed-ENM, we perform a comprehensive mutation-scanning analysis to assess how single-residue substitutions reshape conformational energy landscapes. This framework provides a physically grounded basis for identifying mutation-sensitive residues and substitutions, as well as uncovering global patterns of mutation impact.

## 2. Materials and Methods

### 2.1. Molecular Dynamics simulation dataset

We built two complementary datasets comprising all-atom MD simulation trajectories of proteins with diverse structures and force-field models. The first dataset was generated by performing unbiased MD simulations for five proteins—RBP (271 residues), 5’NTase (525 residues), RNaseIII (438 residues), SERCA (994 residues), and ACLY (4384 residues)—each simulated from pairs of experimentally determined endpoint structures. Simulations were conducted using three different force-field models: AMBER99sb-ildn (26), CHARMM36m (27), and OPLS-AA/L (28). Detailed simulation parameters and system information are provided in Table S1. The second dataset was compiled by collecting MD simulation files of various protein systems indexed in MDverse—a searchable platform designed to aggregate and organize publicly shared MD simulation data (29) (Table S2). These datasets served as the foundation for model development, and their specific roles will be described in the following sections.

### 2.2. PDB structure dataset

We assembled a dataset of PDB structures consisting of 82 endpoint conformations from 41 different proteins, which was used to evaluate the performance of our model. The proteins span a broad range of sizes and structural classes. To define the relative conformational states of each endpoint pair, we assessed structural compactness using the radius of gyration and classified them as either ‘open’ or ‘closed’. A complete list of the proteins their corresponding PDB identifiers, and structural annotations is provided in Table S3.

### 2.3. Internal-coordinate-based, essential-dynamics-refined ENM (ICed-ENM)

The internal coordinate-based, essential dynamics-refined ENM (ICed-ENM) is a refined coarse-grained modeling framework designed to characterize protein structural dynamics and identify mutation-induced changes in the conformational landscape. It simplifies complex protein structures into mass-spring mechanical systems, where selected atoms are treated as mass points, and interatomic interactions as harmonic springs. To interpret molecular geometry in IC space, such as bond lengths, bond angles, and dihedral angles, and to detect geometric changes by mutations, we defined nodes by selecting key heavy atoms: N, Cα, and C from the backbone, and the center of mass of the heavy atoms in each side chain (SCOM) (Fig. 1A). Atomic masses and lumped masses were assigned to the backbone nodes and SCOMs, respectively. Moreover, we assume that the spring constants (k) can be defined by a nonlinear model combining two types of interactions: bonded and non-bonded interactions defined by sequential and spatial distances, respectively. The nonlinear model is given by:

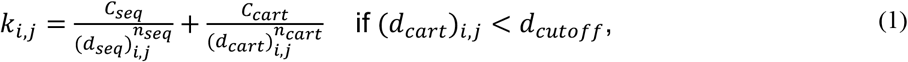

where *k*_*i,j*_ represents the spring constant value between *i*^*th*^ and *j*^*th*^ nodes. (*d*_*seq*_)_*i,j*_ represents the sequential distance between them, indicating how far apart they are in the primary sequence of the molecule. (*d*_*cart*_)_*i,j*_ represents the spatial distance between two atoms in three-dimensional space. *D*_*cutoff*_ is a spatial distance cutoff, defining the range within which neighboring nodes are connected by springs. The parameters *C*_*seq*_ and *C*_*cart*_ represent the prefactors for the sequential and spatial interaction terms, respectively, while *n*_*seq*_ and *n*_*cart*_ denote the corresponding power-law exponents.

**Figure 1.**
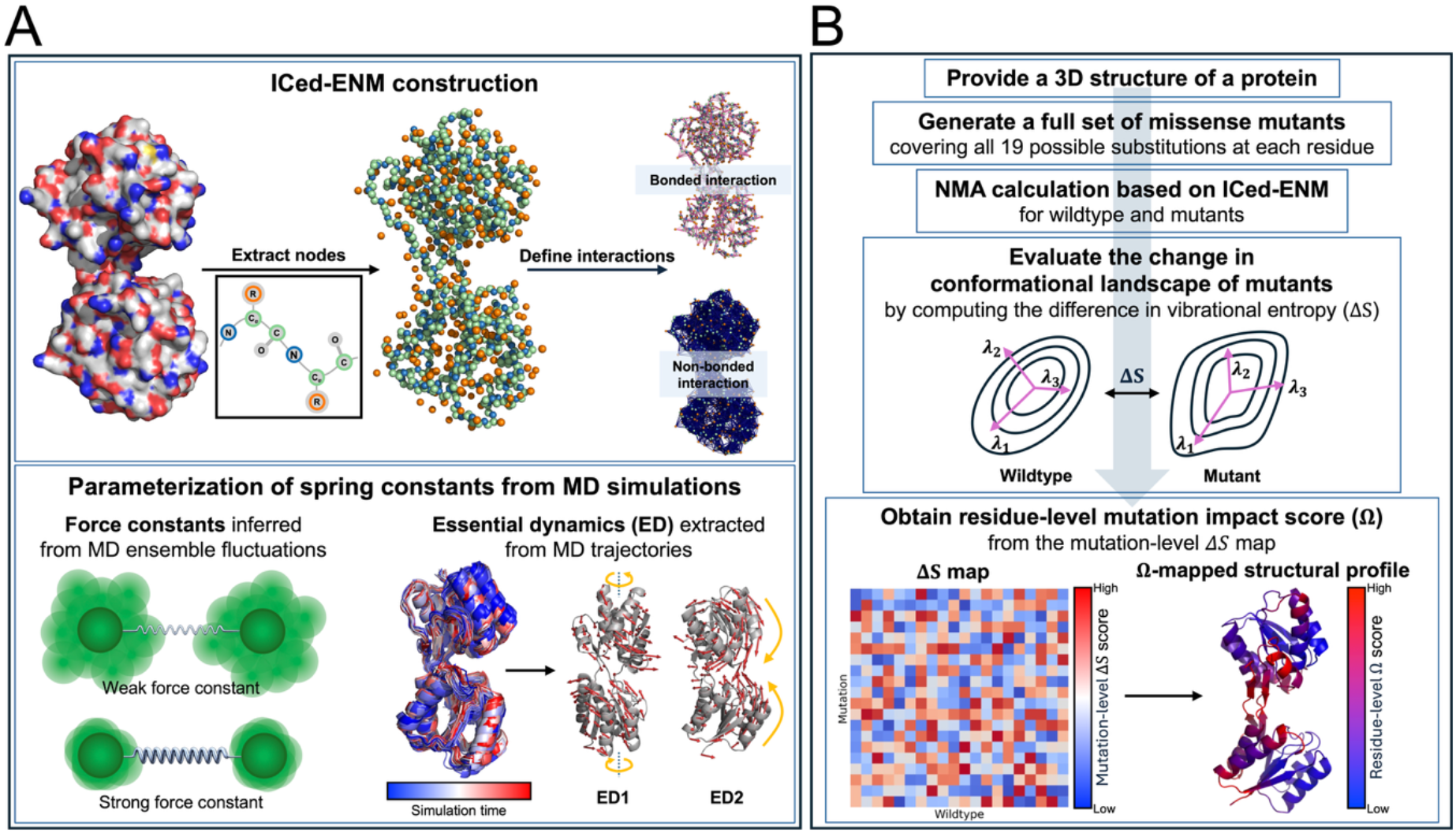
Overview of the internal-coordinate-based, essential-dynamics-refined elastic network model (ICed-ENM) framework. (*A*) Model construction and parameterization. An all-atom structure is coarse-grained into a node-spring network, in which backbone atoms (N, Cα, and C) and the centers of mass of side chains serve as nodes. These nodes are connected by effective springs whose stiffness is defined as the sum of two classes: bonded and non-bonded interactions, defined by sequential and spatial distances, respectively. Model parameterization was performed in two stages using molecular dynamics (MD) simulation data. First, inter-node apparent force constants derived from time-ensemble fluctuations in MD simulations were used to train the model, yielding effective spring constants that reproduce the structural flexibility encoded in the MD ensemble (the lower-left panel). Second, essential dynamics (ED) extracted from MD trajectories were incorporated to refine the lowest normal modes, improving their alignment with the MD-derived essential subspace (the lower-right panel). (*B*) Mutation-impact analysis through the amino-acid scanning approach. In the first step, a full set of single-point missense mutants is generated from the input three-dimensional conformation. Next, normal mode analysis based on ICed-ENM is performed for both the wild-type and the mutants. From these results, the differences in vibrational entropy (*ΔS*) are computed and mapped across amino-acid substitution types and residue positions. By integrating this map with the protein structural information, residue-level mutation impact scores (*Ω*) are finally obtained.

To optimize these parameters for accurately reproducing the physically reliable dynamic behavior, we conducted a two-step parameterization procedure based on the MD simulation datasets described above. In the first step, four scalar parameters of the nonlinear model (Eq. (1)) such as *C*_*seq*_, *C*_*cart*_, *n*_*seq*_, and *n*_*cart*_, were calibrated using apparent force constants (*K*_*app*_) derived from time-ensemble fluctuations in MD trajectories (16), representing effective stiffness between nodes including N, Cα, C, and SCOM:

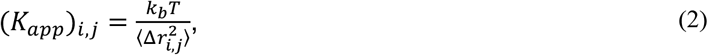

where *k*_*b*_ is the Boltzmann constant and *T* is the absolute temperature (303.15 K in this study). The term 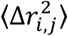 represents the time ensemble average of the squared fluctuations in the distance between *i*^*th*^ and *j*^*th*^ atoms from their mean distance over the trajectory. The nonlinear model training was carried out using 10-fold cross-validation on the local dataset (18,480,049 samples with 26 MD trajectories) (30). Fig. S1*A* shows the actual profile of *K*_*app*_ as a function of *d*_*seq*_ and *d*_*cart*_ for the dataset. The model performance was then evaluated on the MDverse dataset (13,807,275 samples with 21 trajectories). This procedure yielded the following optimal parameter values: *C*_*seq*_: 286.4, *C*_*cart*_: 147.6, *n*_*seq*_: 1.92, and *n*_*cart*_: 1.66.

In the second step, we conducted another parameterization study for *d*_*cutoff*_. To find the optimum cutoff value that best reproduces the energy landscape sampled by MD simulations, the similarity between the subspaces spanned by normal modes of ICed-ENM (with the already obtained optimum parameters) and those spanned by the essential dynamics (ED) from the total MD dataset combining the local and MDverse datasets was quantified using the root-mean-square inner product (RMSIP), which measures the overlap between two mode subspaces. Fig. S1*B* shows the mean and median RMSIP values across the MD dataset for the lowest 3, 5, and 10 modes as a function of *d*_*cutoff*_. Based on these results, we selected the optimum *d*_*cutoff*_ (set to be 8 Å), corresponding to the point at which our model achieves the highest performance in reproducing the flexibility of the all-atom model of MD simulations.

### 2.4. Normal mode analysis (NMA) based on ICed-ENM

Normal mode analysis (NMA) based on ENMs is a widely used computational technique for characterizing the vibrational and dynamic properties of proteins at equilibrium (17). It employs a harmonic approximation of the potential energy surface and solves an eigenvalue problem derived from the Hessian matrix, which contains the second derivatives of the potential energy with respect to atomic displacements (optionally incorporating a mass matrix to account for atomic weights). Each normal mode consists of an eigenvector and its corresponding eigenvalue, representing the direction and frequency of the intrinsic motion, respectively. In particular, the low-frequency normal modes are often associated with large-scale conformational changes that are functionally relevant.

In this study, we performed NMA using ICed-ENM, which characterizes structural dynamics based solely on backbone dihedral angles (such as *ϕ* and *ψ* angles). To carry out the NMA, we first expressed both the potential and the kinetic energy functions in the IC space, enabling the construction of the Hessian and global mass matrices, corresponding to the second derivatives of potential and kinetic energies, respectively (20, 31). Let **θ** = (*θ*_1_, *θ*_2_, ⋯, *θ*_*n*_)^*T*^ denote the column vector of ICs (i.e., *ϕ* and *ψ* angles). The Hessian matrix (**H**) in the IC space is given by:

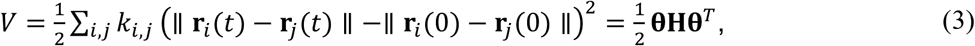

where,

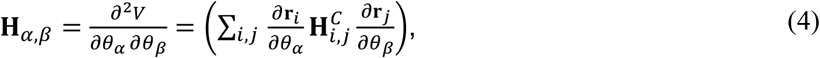

*i* and *j* (*α* and *β*) are the indices in CC (IC) space, **r**_*i*_(*t*) is the position vector of the *i*^*th*^ atom at time *t*, and 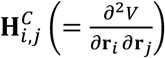 is the Hessian matrix in Cartesian coordinates. Similarly, the global mass matrix (**M**) is obtained from the kinetic energy:

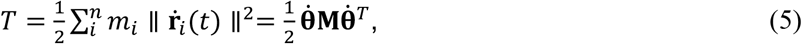

where *m*_*i*_ is the mass of the *i*^*th*^ atom and

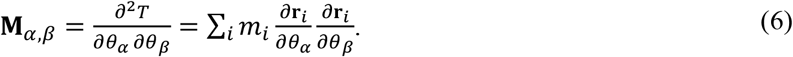

Finally, the generalized eigenvalue problem **Hv** = *λ***Mv** is solved to obtain the eigenvalues (*λ*) and the eigenvectors (**v**), which represent the mode frequencies and their corresponding displacement patterns, respectively.

### 2.5. Mutation-impact analysis through an amino-acid scanning approach

As noted above, ICed-ENM represents not only the positions of backbone atoms but also approximate side-chain geometry, enabling the model to account for side-chain-mediated effects. Building on this capability, we applied the model to evaluate the impact of missense mutations on the conformational energy landscape. Fig. 1*B* illustrates the overall workflow of the application. Starting from an input three-dimensional conformation (typically a PDB structure), all possible single-point missense mutants are generated by substituting each residue with the 19 other standard amino acids using Modeller 10.7 (32). For the wildtype structure (WT) and all possible mutants, ICed-ENM-based NMA is performed, and the changes in the conformational energy landscape induced by the mutants are quantified by computing the absolute differences in vibrational entropy (*ΔS*) derived from the resulting normal modes. The *ΔS* values serve as a mutation-level metric quantifying how individual substitutions reshape the conformational energy landscape of the WT. To derive the residue-level mutation impact score (*Ω*), additional post-processing steps are applied. First, for each residue, the maximum *ΔS* among the 19 substitutions is selected to capture position-specific mutation sensitivity. To incorporate local structural context, a neighboring list is defined for each residue based on C*α* coordinates using a 15 Å distance cutoff. Gaussian kernel smoothing is then applied to the selected *ΔS* values within this spatially defined neighborhood, allowing local mutation sensitivities to be refined based on structural proximity. To avoid artifacts arising from weakly connected regions, residues with fewer than six C*α* neighbors within an 8 Å distance cutoff are excluded from the Gaussian smoothing procedure. This strategy mitigates artifacts arising from weakly connected or terminal regions. Finally, the resulting *Ω* profile is mapped onto the protein structure.

### 2.6. Root-mean-square inner product (RMSIP)

The root-mean-square inner product (RMSIP) quantifies the similarity between two sets of normal modes (33). A higher RMSIP value indicates greater overlap between the corresponding mode subspaces. In this study, the RMSIP was used to assess the similarity between mode subspaces obtained from ENM-based NMA and ED from MD trajectories. The equation is given by:

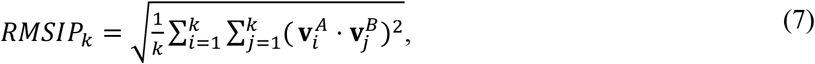

where *k* is the number of modes considered in the comparison, and 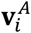 and 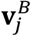 denote the *i*^*th*^ and *j*^*th*^ normalized eigenvectors from the mode sets *A* and *B*, respectively.

### 2.7. Essential dynamics (ED) via principal component analysis (PCA)

Essential dynamics (ED), a principal application of principal component analysis (PCA), is a powerful technique in MD simulations to characterize the dominant modes of atomic motion by examining the covariance matrix of atomic positional fluctuations over time (34). It involves computing the eigenvectors and eigenvalues of the covariance matrix, where each eigenvector represents a principal direction of motion, and its corresponding eigenvalue indicates the amplitude of fluctuation along the direction. This reduction in dimensionality highlights a small number of collective modes that effectively capture the essential dynamic behavior of the system. In this study, ED derived from MD trajectories was used to guide the parameterization of our proposed model.

### 2.8. Difference in vibrational entropy

The difference in the vibrational entropy along the conformational landscape effectively captures the variation in structural flexibility and stability between the two conformations (21, 35). In this study, we computed this difference between the WT and mutant systems to evaluate the sensitivity of our proposed model to point mutations. The entropy change from conformation A to conformation B (*ΔS*_*A*→*B*_) is calculated as:

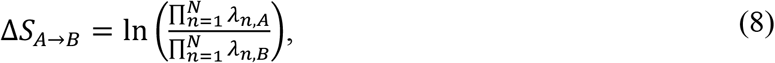

where *λ*_*n,i*_ are the *n*^*th*^ eigenvalue from NMA calculation for conformation *i*. When *ΔS*_*A*→*B*_ < 0, it means conformation *B* is less flexible than conformation *A* (and vice versa). Because we are interested in the magnitude rather than the direction of the change, the absolute value of *ΔS* is considered.

### 2.9. Cumulative square overlap (CSO)

The cumulative square overlap (CSO) is a metric used to evaluate the similarity between a single vector and a subspace spanned by a set of normal mode vectors (36). A CSO score of 1 implies that the selected subspace can fully represent the transition vector. In this study, CSO was used to assess how well a normal mode subspace captures the transition vector between two-endpoint conformations. The CSO score of the subspace composed of the first *k* normal mode vectors is calculated as:

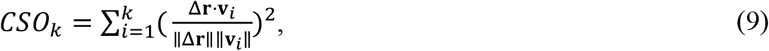

where Δ**r** is the transition vector, and **v**_*i*_ is the *i*^*th*^ normal mode vector.

### 2.10. Potential of mean force (PMF) in the principal component (PC) subspace

The potential of mean force (PMF) provides a free-energy profile projected onto selected collective variables by integrating out all remaining atomic degrees of freedom (37). In this study, conformations sampled from MD trajectories were projected onto the first two PCs derived from the experimental PDB ensembles to characterize mutation-induced reshaping of the conformational energy landscape. The PMF, expressed as a function of the collective variables *s*_1_ and *s*_2_ (corresponding to PC1 and PC2), was computed using the Boltzmann relation:

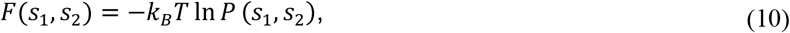

where ln *P* (*s*_1_, *s*_2_) is the probability density of conformations located at the position (*s*_1_, *s*_2_) in the PC1–PC2 subspace. The probability density was estimated from merged trajectories of three independent 100-*ns* MD replicas for each system. The resulting PMF was shifted so that its minimum value is zero, enabling direct comparison of relative free-energy differences between the WT and mutant systems.

## 3. Results

### 3.1. Experimental validity of normal mode shapes

To assess the quality of ICed-ENM, we did a benchmark test with different ENM methods: iMOD (edENM force-field with ICs) (38), edENM (with CCs) (16), and ANM (standard unit spring constant with Cartesian coordinates) (17). They are characterized by distinct force-field models and coordinate systems. For our PDB dataset consisting of 41 protein pairs (a total of 82 structures termed as ‘A’), allocated into two groups: open state group (‘O’) and closed state group (‘C’), we performed normal mode analysis (NMA) based on the four ENM models. We first compared them with the information on experimental transitions defined by two endpoint PDB structures, to evaluate how well the models describe the experimental conformational changes. Table 1 reports average overlaps across all protein groups.

**Table 1.**
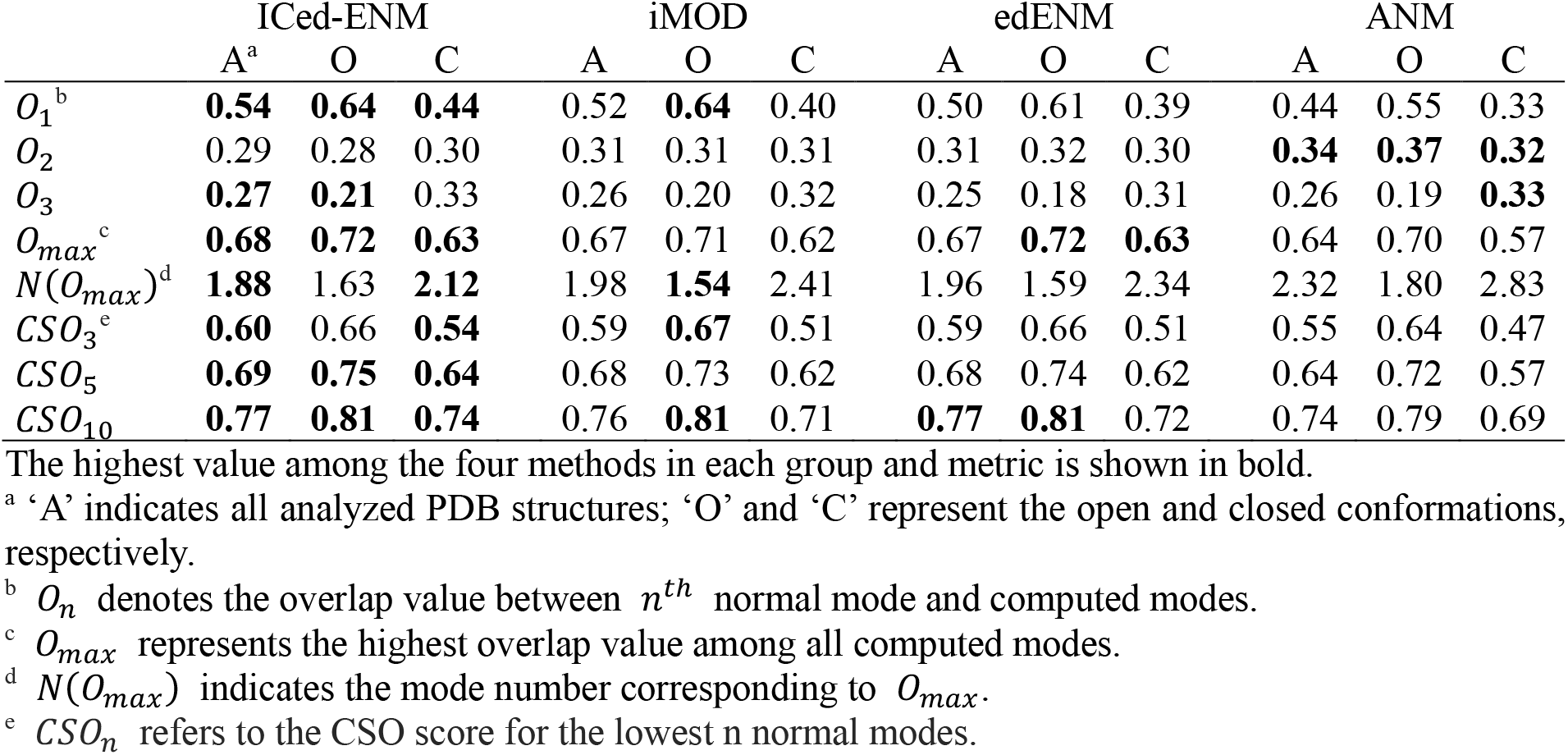
Overlap analysis between computed normal modes and experimental transition vectors.

Across all methods, the overlap of the first normal mode (*O*_1_) with the experimental transition vector exhibits the highest scores, suggesting that this mode correlates most significantly with the conformational transition. All methods showed higher *O*_1_ scores for the ‘O’ group than for the ‘C’ group, indicating that NMA more effectively captures conformational transitions when using open conformations as input. Among the methods, ICed-ENM produced the highest *O*_1_ scores across all structural groups: 0.54 for ‘A’, 0.64 for ‘O’, and 0.44 for ‘C’ whereas ANM yielded the lowest, with scores of 0.44, 0.55, and 0.33, respectively. In contrast, the overlap score for the second normal mode (*O*_2_) showed an inverse trend: ICed-ENM yielded the lowest *O*_2_ scores (0.29, 0.28, 0.30), while ANM produced the highest (0.34, 0.37, 0.32). This pattern indicates that ICed-ENM concentrates most of the transition-related motions into the first mode, thereby reducing the contributions from higher modes. Otherwise, the transition-related information becomes more diffusely distributed, as is often observed with other methods. In addition, the maximum overlap scores (*O*_*max*_) and their corresponding mode indices (*N*(*O*_*max*_)) were evaluated to assess model performances. ICed-ENM consistently achieved the highest Omax scores coupled with the lowest *N*(*O*_*max*_), indicating that it captures the most relevant dynamic information at lower mode indices compared to the other methods. Additionally, the CSO score was computed to assess how well a subspace composed of lower modes collectively spans the transition vector. It was evaluated using the lowest 3, 5, and 10 normal modes (*CSO*_3_, *CSO*_5_, and *CSO*_10_). ICed-ENM tended to produce higher CSO scores across most subsets and structural groups, suggesting that it effectively captures transition-relevant motions within a small number of low-frequency modes.

### 3.2. Stereochemical validity of normal mode shapes

As mentioned above, ICed-ENM utilizes the IC space for analyzing protein dynamics. The space inherently reflects the structural characteristics of molecules, which enables NMA to produce more chemically realistic dynamics by efficiently constraining redundant degrees of freedom (e.g., bond-length and bond-angle), while maintaining flexibility in key dihedral angles (e.g., *ϕ* and *ψ*). To assess the impact of our model on the resulting normal modes, we analyzed several metrics related to the stereochemical stability of protein structures and performed the same benchmark comparison with conventional ENM methods for our PDB dataset (Fig. 2).

**Figure 2.**
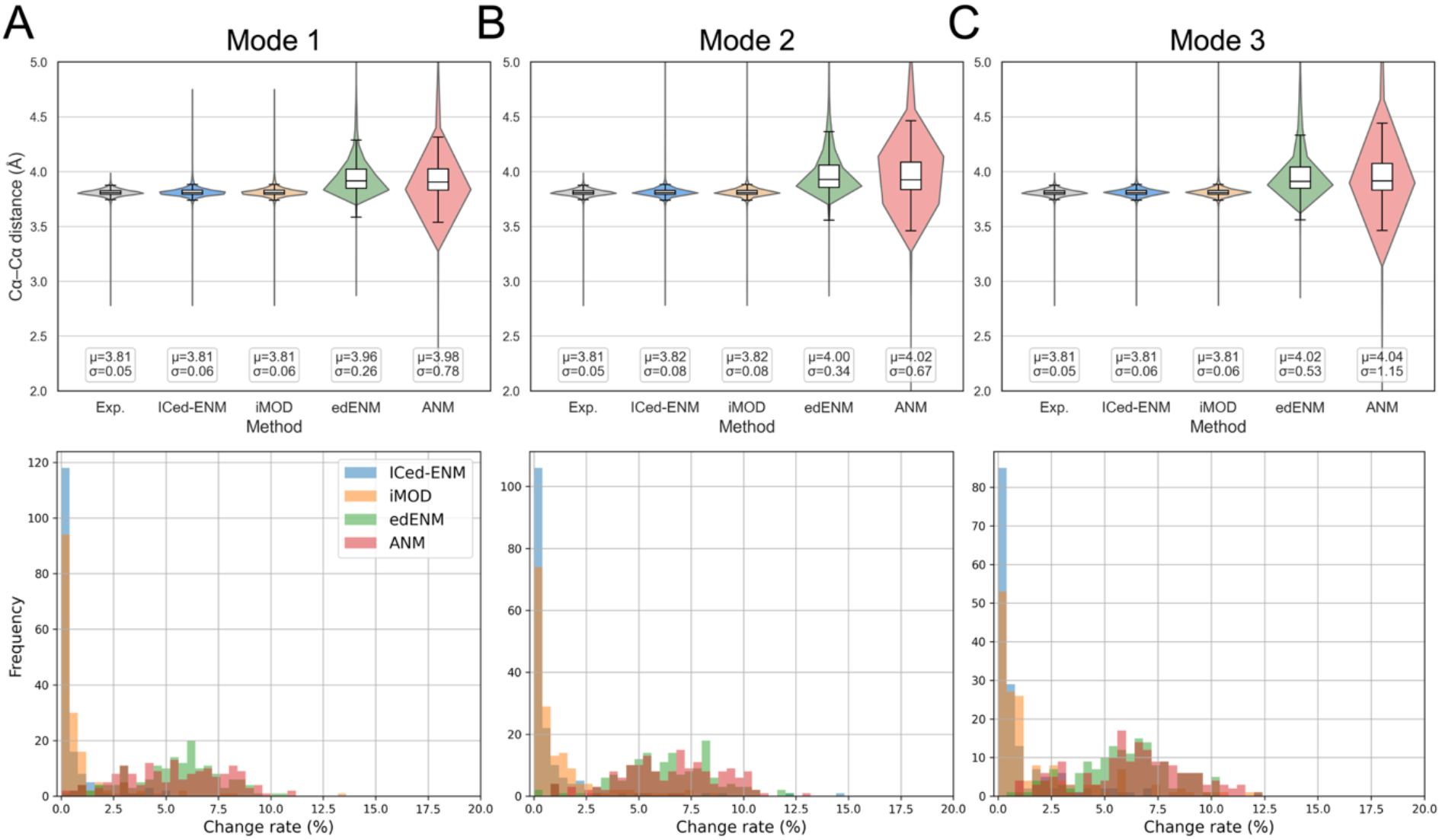
Evaluation of stereochemical stability of low-frequency normal modes. (*A*–*C*) Each column corresponds to one of the first three low-frequency normal modes: Mode 1 (*A*), Mode 2 (*B*), and Mode 3 (*C*). The top row shows violin plots of sequential C*α*–C*α* distance distributions for the experimental structures (Exp., gray) and structures deformed along the first three normal modes (Modes 1–3) using different ENM models (ICed-ENM, iMOD, edENM, and ANM; blue, orange, green, and red, respectively). The bottom row shows histograms of bond-count change rates for neighboring interactions between C*α*s. The change rate was calculated by comparing the number of local bonds of the deformed structures with those in the corresponding experimental structures, using a cutoff distance of 8 Å.

Deformed structures were obtained using the normal modes obtained from each ENM model. For each mode, two deformations were produced by displacing the structure in the positive and negative directions of the mode vector, with a displacement magnitude of 5 Å from the original structure. For all deformed structures, we computed the distances between the sequentially connected C*α* atoms (C*α*–C*α* distances). Since the C*α*–C*α* distance is expected to remain close to ~3.8 Å to preserve chemically reasonable backbone geometry, this analysis provides a measure of how well the backbone framework is conserved along the mode shapes. The top panels of Fig. 2 shows the distributions of C*α*–C*α* distances using violin plots. Aside from a small number of outliers, the distributions for structures deformed using the ENMs based on IC space (ICed-ENM and iMOD) are tightly concentrated around ~3.8 Å, similar to those of the experimental structures, indicating the backbone geometry is largely preserved along the mode shapes. In contrast, structures deformed by the other two ENMs based on CC space (edENM and ANM) exhibit more dispersed distance distributions, implying reduced stereochemical stability in the deformed conformations.

Furthermore, we evaluated the overall stability of the deformed conformations using a complementary, network-level metric by quantifying changes in the total number of neighboring C*α* connections. Specifically, we computed the change rate in the total number of the bonds in the C*α* networks by comparing the deformed structures to the original experimental structures. The networks are constructed using a cutoff distance of 8 Å. The bottom panels of Fig. 2 show the distributions of bond-count change rates for each normal mode. Consistent with the results of the C*α*–C*α* distance analysis, the IC-based models exhibit lower change rates than CC-based models. Particularly, ICed-ENM yields the most stable and conserved mode shapes in the conformational landscape.

### 3.3. Ability to capture mutation-induced perturbations in the conformational energy landscape

To assess whether ICed-ENM reliably captures mutation-induced perturbations of the conformational energy landscape, we performed our mutation-impact analysis on two well-characterized systems: E. coli ribose-binding protein (RBP) and E. coli 5’-nucleotidase (5’NTase). Mutation targets were selected based on residue-level *Ω* profiles, with high-*Ω* residues defined as hot spots. These hot-spot targets were further refined to span diverse structural contexts, including amino-acid type, solvent-accessible surface area (SASA) computed using FreeSASA (39), and secondary-structure class (helix, strand, and coil) assigned using DSSP (40). Low-*Ω* residues (cold spots) corresponding to the selected hot spots were then chosen as controls, considering both structural context and physicochemical characteristics to enable balanced comparison of mutation effects. To validate the predicted mutation sensitivity, all-atom MD systems were constructed using the CHARMM-GUI web interface (41), where single-point alanine substitutions were introduced at the selected hot- and cold-spot positions. MD simulations were performed using three independent 100-*ns* replicas with the CHARMM36m force field. To quantify the impact of each mutation, the resulting trajectories were projected onto the PC1–PC2 subspace defined by experimental PDB ensembles (11 structures for RBP and 13 for 5’NTase; see Table S3). Conformational free-energy landscapes were then estimated as the potential of mean force (PMF) in this PC subspace.

RBP, a member of the periplasmic binding protein family, undergoes a hinge-like motion that enables ribose binding and plays a key role in ligand transport (42). For the apo open conformation corresponding to chain A of PDB entry 1URP (denoted as 1URP:A), we computed the residue-level *Ω* profile and selected two pairs of hot- and cold-spot residues: (i) Q235 (SASA: 33.7%) and its low-*Ω* counterpart Q184 (44.3%), and (ii) E255 (54.9%) and E122 (66.1%) (left structural model in Fig. 3*A*). Q235 is located in a loop within the canonical hinge region mediating inter-domain motion, and Q184 occupies a comparable loop environment. E255 resides in a solvent-exposed loop of the N-terminal domain, and E122 is positioned in a similar solvent-exposed loop region. Projection of the WT and mutant trajectories onto the PC1–PC2 subspace reveals mutation-dependent reshaping of the conformational free-energy landscape (landscape plots in Fig. 3*A*). The WT system exhibits a dominant basin centered near the reference conformation, and the cold-spot MUT systems (Q184A and E122A) display landscapes similar to that of WT. In contrast, Q235A retains the original while clearly populating a secondary basin shifted toward positive PC1 and PC2 values, suggesting altered sampling along the inter-domain hinge mode. Similarly, E255A exhibits a notably more diffuse distribution across the PC subspace, forming a distinct auxiliary basin that indicates increased conformational heterogeneity.

**Figure 3.**
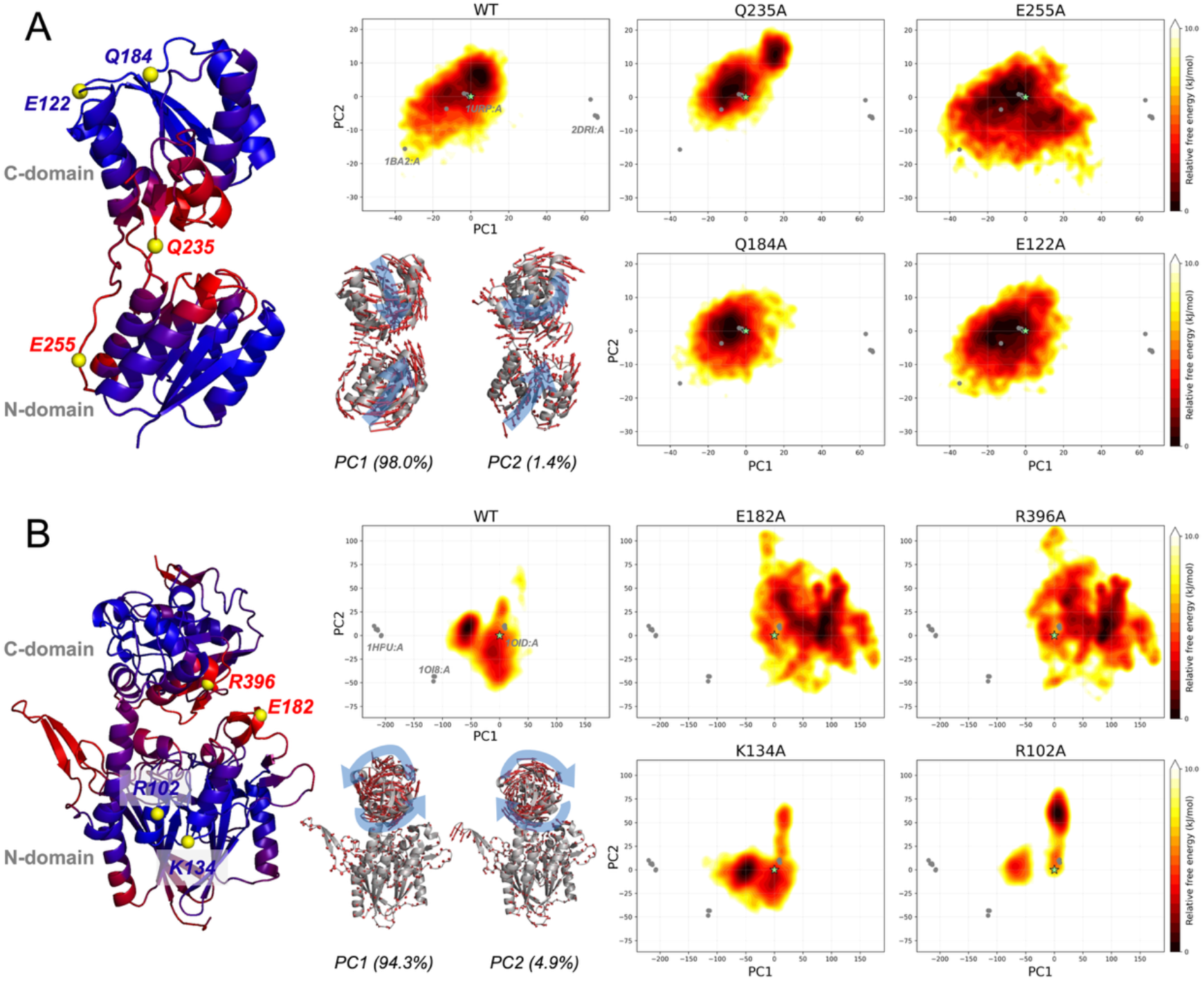
Mutation-induced reshaping of conformational free-energy landscapes projected onto experimental principal component (PC1–PC2) subspaces. The left panels display the *Ω* profiles mapped onto the three-dimensional structures of 1URP:A in RBP (*A*) and 1OID:A in 5’NTase (*B*). Based on these profiles, two representative high-impact residues (hot spots) were selected for alanine substitution, together with structurally comparable low-score residues (cold spots) as counterparts for control comparisons. The small structural panels illustrate the dynamic shapes of PC1 and PC2 derived from the experimental ensembles. Conformational free-energy landscapes were constructed from three independent 100-*ns* all-atom MD replicas for WT and each mutant and represented as potentials of mean force (PMF), with the minimum free energy shifted to zero. Gray points denote projections of experimental PDB structures, and the light-green star marks the reference PDB structure. (*A*) In 1URP:A, Q235 (canonical hinge loop) and E255 (solvent-exposed loop) were selected as hot spots, with Q184 and E122 as counterparts. PC1 describes the dominant inter-domain hinge motion between the N- and C-terminal domains, while PC2 represents a minor twisting motion around the hinge axis. (*B*) In 1OID:A, R396 and E182, located at inter-domain contact regions, were selected as hot spots, with R102 and K134 as counterparts. PC1 and PC2 represent distinct rotational motions of the C-terminal domain relative to the N-terminal domain.

5’NTase, a periplasmic enzyme involved in nucleotide metabolism, consists of N- and C-terminal domains connected by an *α* helix that enable inter-domain rearrangements during catalysis (43). While 1OID:A represents an apo open conformation, it contains engineered cysteine substitutions (S228C and P513C) that form a disulfide bridge. Therefore, these residues were reverted to their native identities, and the resulting structure was used as the WT reference. Following the same protocol, we calculated the residue-level *Ω* profile and selected two pairs of hot- and cold-spot residues: (i) E182 (SASA: 84.9%) and its low-*Ω* counterpart K134 (85.7%), and (ii) R396 (28.7%) and R102 (33.1%) (left structural model in Fig. 3*B*). E182 is located in a solvent-exposed loop region of the N-terminal at the inter-domain interface, and K134 occupies a comparable solvent-exposed loop environment. Although E182 and K134 differ in amino-acid identity (acidic versus basic), both are charged residues, allowing comparison within a similar electrostatic context. R396 resides in an α-helical segment within the C-terminal domain at the inter-domain interface, and R102 is positioned in a distinct *α*-helical region. The conformational free-energy landscapes, mapped onto the PC1–PC2 subspace, exhibit distinct mutation-induced shifts (landscape plots in Fig. 3*B*). The WT system is characterized by a primary basin shifted toward negative PC1 values relative to the reference, with a secondary basin in its vicinity. The cold-spot mutant systems largely preserve this landscape; although R102A samples an additional basin toward positive PC2 values, the overall distribution remains comparable to that of WT. In contrast, the hot-spot mutants (E182A and R396A) radically redistribute conformational populations, occupying vast new areas of the PC subspace and indicating a pronounced alteration of the underlying energy landscape.

### 3.4. Analysis of global mutation-impact patterns

To statistically characterize the mutation-induced perturbations, we applied the ICed-ENM-based mutation-impact analysis to our PDB dataset. For all PDBs except ACLY having missing regions, we computed mutation-level *ΔS* scores and residue-level *Ω* scores and yielded a large-scale mutation-impact dataset, enabling global statistical analyses across diverse structural and dynamic contexts. First, we evaluated whether mutation-impact scores are associated with structural deviations of mutant structures relative to WT, as quantified by RMSD (Fig. 4*A*). To the end, mutations were grouped into the top 10% *ΔS* group and the remaining 90%. Despite pronounced differences in *ΔS*, the RMSD distributions of the two groups are highly overlapping. Importantly, the inset displaying the raw RMSD distribution reveals that absolute structural deviations remain exceedingly small, with a median RMSD of ~ 0.006 Å. Together, these results demonstrate that mutation impacts quantified by our approach are not trivially explained by the magnitude of structural deviation.

**Figure 4.**
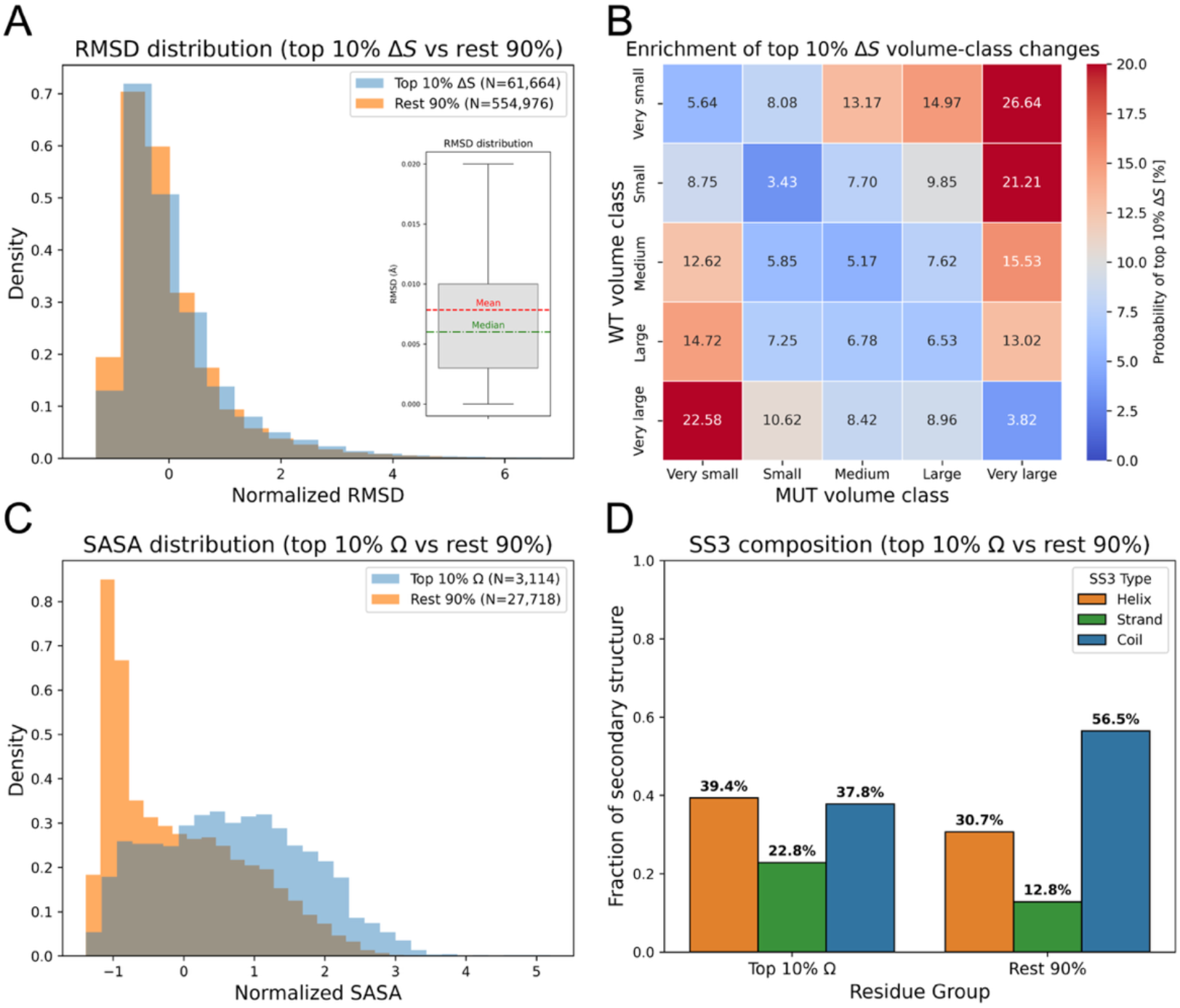
Statistical analysis for the mutation-level *ΔS* score (*A* and *B*) and the residue-level *Ω* score (*C* and *D*). (*A*) Distribution of mutation-level RMSD of mutants relative to the WT for residues in the top 10% *ΔS* group versus the remaining 90%. RMSD values were Z-score-normalized per structure. The inset shows the raw RMSD distribution across all mutations. (*B*) Probability of belonging to the top 10% *ΔS* group as a function of the WT and mutant side-chain volume classes. The color scale is centered at the global expectation of 10%, such that warmer colors indicate volume-class transitions that generate high *ΔS* scores more frequently than expected by chance. (*C*) Distribution of residue-level solvent-accessible surface area (SASA) for residues ranked within the top 10% of *Ω* versus the remaining 90%. SASA values were Z-score-normalized per structure. (*D*) Secondary-structure composition of residues in the top 10% *Ω* group compared with the remainder, based on a three-state (SS3) classification (helix, strand, and coil). Bars represent the fraction of residues within each group.

Next, we investigated the effects of side-chain volume changes induced by single-point mutations on *ΔS* score (Fig. 4*B*). To facilitate interpretation, amino-acid side-chain volumes were grouped into five classes (44), and all possible WT-to-mutant class transitions were systematically evaluated. The heatmap reports the probability (%) that a given volume-class transition produces high *ΔS* values, defined as the top 10% of *ΔS* within each structure; thus, 10.0% corresponds to the background expectation. A prominent pattern emerges: large-magnitude volume changes are generally associated with elevated probabilities of high *ΔS* responses. Notably, transitions from smaller to large volume classes exhibit stronger enrichment than transitions in the reverse direction. In particular, mutations originating from very large residues toward smaller classes, except for transitions to the very small group, yield probabilities close to or below the background level. This analysis highlights systematic biases in the occurrence of large vibrational-entropy changes that depend on the magnitude and direction of side-chain volume perturbations.

Aside from the mutation-level analysis, we further analyzed residue-level *Ω* scores. Fig. 4*C* shows the distribution of residue-level SASA for the high-*Ω* group (3,113 residues), defined as the top 10% of *Ω* within each structure, and the remaining residues (27,718 residues). A clear difference is observed between the two groups. The remaining residues show a strong enrichment at low SASA values, consistent with the intrinsic geometric packing of protein structures, where buried residues predominate. In contrast, the high-*Ω* group exhibits a broader and more gradually distributed SASA profile. Rather than simply reflecting the baseline bias toward buried positions, high-*Ω* residues extend across a wide range of solvent accessibility, indicating that mutation sensitivity is not determined solely by solvent exposure. This observation is further reflected in the distribution of residue-level number of contacts (NOC) for the *Ω* group (Fig. S2). The high-Ω group shows a modest shift toward lower NOC values compared to the remaining group, consistent with their increased representation in more solvent-exposed regions. This analysis supports the SASA findings from a structural packing perspective.

To further characterize the structural context of high-*Ω* residues, we examined their secondary-structure distribution using a three-state (SS3) classification (Fig. 4*D*). The SS3 scheme groups residues into three structural states: helix (*α*-helical regions), strand (*β*-strand segments), and coil (loops or irregular regions). Compared to the baseline distribution of the remaining residues, helices and strands show relative enrichment in the high-*Ω* group, whereas coils are proportionally reduced. In absolute terms, high-*Ω* residues are most frequently observed in helices (39.4%), followed by coils (37.8%), while strand regions comprise a smaller fraction (22.8%) overall.

## 4. Discussion

In this study, we introduced ICed-ENM, a framework designed not only to capture functionally relevant collective motions of proteins but also to sensitively detect mutation impacts on conformational energy landscapes. Benchmark analyses demonstrate that ICed-ENM generates normal modes that show both improved agreement with experimentally observed transition directions and enhanced stereochemical reliability. This improvement likely arises from the IC representation, which constrains bond-length and bond-angle fluctuations while preserving physically meaningful torsional motions. Although the overall improvement relative to iMOD is moderate, as both methods share the fundamental advantages of IC, this analysis is essential to establish a stable, physically grounded reference for detecting subtle mutation-induced perturbations without artificially amplifying global motions.

The mutation-scanning analysis based on ICed-ENM demonstrates its applicability in identifying mutation-sensitive regions within protein conformational landscapes. The mutation-impact scores employed here are derived from the difference in vibrational entropy between the WT and mutant systems, thereby providing a quantitative measure of mutation-induced redistribution within the conformational energy landscapes based on structure-encoded intrinsic dynamics. To validate this approach, residues predicted as mutation hot spots were evaluated by comparing changes in free-energy landscapes derived from all-atom MD simulations of alanine mutants at the hot spots with those at structurally and chemically matched cold spots. Notably, E255A in 1URP:A, located within a solvent-exposed loop region, was predicted to exert a considerable impact on functionally relevant inter-domain hinge motions (Fig. 3*A*). These findings indicate that the framework not only captures intuitively critical regions, such as inter-domain hinge regions or domain interfaces, but also identifies less obvious positions whose perturbations propagate beyond their local structural environments. The ability to detect mutation effects consistent with long-range allosteric coupling represents a key strength of the present approach, particularly given that mutations in loops or other distal, apparently benign regions are notoriously difficult to interpret based solely on static structural or energetic considerations (5, 6).

Extending this analysis beyond individual case studies, we examined the global patterns of mutation hot spots identified by our mutation-scanning framework across our curated PDB dataset (Fig. 4). To evaluate the broader relevance and potential significance of these patterns, we compared our score trends with those observed in large-scale public mutation datasets. First, we investigated this relationship from a physicochemical perspective. Our results already exhibit pronounced sensitivity to amino-acid volume changes, indicating that steric perturbations are effectively captured within the elastic network representation (Fig. 4*B*). To gain complementary insight into this relationship, we examined substitution-specific enrichment patterns across four major physicochemical classes (hydrophobic, polar uncharged, negative, and positive) (Fig. S3*A*). A clear trend emerged: substitutions that conserve physicochemical properties generally displayed enrichment values below the baseline, suggesting reduced mutation impact. This pattern mirrors observations from human disease mutation analyses, including studies based on the HumVar dataset (45, 46) and comparative analyses of disease-associated versus polymorphic mutations (47), where conservative substitutions are generally underrepresented among pathogenic variants. In contrast, substitutions involving transitions from charged residues (either positive or negative) to hydrophobic residues exhibited markedly elevated enrichment scores in our framework. These findings can be partially explained by changes in salt bridges that has a significant impact on protein dynamics and is known to be a factor for disease-related mutations (48, 49). Although ENM-based models do not explicitly encode specific chemical bonds or charges, our results suggest that the resulting perturbations in the elastic interaction network are translated into detectable changes in conformational dynamics. In this sense, network-level mechanical changes appear to serve as an indirect proxy for chemically disruptive events.

A more fine-grained analysis at the level of individual amino-acid identities highlights residue-specific sensitivities (Fig. S3 *B* and *C*). In particular, substitutions involving arginine exhibit systematically elevated probabilities of belonging to the high *ΔS* category, even when the associated volume change is modest. This pattern is consistent with prior observations that arginine-related mutations are disproportionately associated with pathogenic propensity (45, 50). Given that Arginine residues frequently participate in salt bridges and extensive hydrogen-bond networks, these findings also suggest that our framework sensitively captures the disruption of such electrostatic interactions induced by arginine substitutions, extending beyond simple steric effects. In addition to arginine, glycine- and proline-related substitutions also display elevated mutation-impact probabilities, even within transitions typically considered conservative in terms of side-chain volume. Given that glycine and proline uniquely regulate backbone flexibility, these observations indicate that the framework captures residue-type-specific backbone effects that are distinct from simple volume or generic chemical-class changes.

From a structural-context perspective, several aspects of our mutation-impact patterns can also be interpreted in light of trends reported in public mutation datasets. Previous studies have reported only marginal differences in SS3 fractions between disease-causing and polymorphic mutations, with coil regions typically being most frequent, followed by helices and strands (45, 48). This general distribution closely matches the baseline proportions observed in our dataset, as reflected in the remaining 90% residue set. In contrast, the high-*Ω* group exhibits modest deviations from this baseline SS3 distribution, which suggests that mutation pathogenicity is unlikely to be determined by secondary-structure identity alone but rather other structural or physicochemical determinants. In terms of solvent accessibility, previous studies have consistently shown that disease-causing mutations are enriched in buried or partially exposed regions compared with polymorphic mutations (45, 51–53). Our results similarly indicate that mutation-sensitive residues are preferentially located in buried or partially exposed environments relative to the remaining residue set. Such residues are more likely to participate in dense interaction networks involving hydrogen bonds, salt bridges, and packing contacts. Because the ENM framework encodes mutation impact as perturbations in mechanical coupling between residues, these findings provide a mechanistic rationale for why mutation pathogenicity may be, at least in part, embedded in the topology of the structure-derived interaction network.

Important limitations remain in our approach. First, ENM is known to exhibit non-physical fluctuations in poorly connected or terminal residues (54), which may lead to an artificial inflation of predicted mutation impacts in these regions. Although we mitigated this issue by filtering weakly connected residues during the *Ω* score calculation, this intrinsic limitation of coarse-grained elastic models cannot be entirely eliminated. Consequently, some residual noise may still be present in our mutation-impact estimates. Second, our framework does not explicitly account for fine-grained chemical properties when quantifying mutation-induced perturbations in conformational dynamics. For example, cysteine mutations are frequently associated with elevated disease risk due to their critical roles in disulfide bond formation and metal coordination (45, 55, 56). In our analysis, however, the mutation impact of cysteine substitutions is comparatively underestimated, which reflects the fact that coarse-grained ENM-based models primarily represent mechanical interaction networks rather than specific chemical functionalities. Third, our framework does not directly evaluate mutation-induced changes in folding free energy, which have been widely used to assess thermodynamic destabilization in disease-associated variants (4, 57, 58). As a result, mutations whose primary impacts arise from large-scale folding destabilization may not be fully captured by our mutation sensitivity metric.

Despite these limitations, our results demonstrate that the proposed framework captures mutation hot spots from the perspective of conformational dynamics, as supported by all-atom MD simulations and broad qualitative consistency between our scores and experimentally characterized mutation propensities. These findings suggest that a substantial component of mutation pathogenicity is encoded in structure-derived dynamics and mechanical interaction networks. Importantly, our framework is deliberately confined to this dynamic layer of mutation impact. It does not aim to encompass the full spectrum of pathogenic mechanisms, which arise from a complex interplay of sequence, structure, dynamics and cellular context. Rather, our approach provides a systematic pipeline for identifying candidate mutation hot spots specifically from the perspective of conformational fluctuations. In this sense, the framework may serve as a complementary and mechanistically interpretable tool that elucidates how local perturbations propagate through protein interaction networks and reshape conformational energy landscapes.

## 5. Author Contribution

B.H.L. and L.O. designed the research; B.H.L. performed the research; B.H.L., D.S., and S.P. analyzed data; B.H.L. wrote the paper; and L.O. supervised the research.

## 6. Acknowledgements

L.O. acknowledges financial support from Cancerfonden Junior Investigator Award (CF 21 0305 JIA) and Project Grants (CF 21 1471 Pj, CF 24 3801 Pj) as well as Vetenskapsrådet Starting Grant (VR 2021-02248). D.S. acknowledges financial support from Cancerfonden Postdoctoral Fellowship (CF 24 0908 PT). S.P. acknowledges financial support from a Karolinska Interdisciplinary Research Incubator (KIRI) postdoctoral grant.

