## Supplementary Information for "Mutation-induced reshaping of protein conformational dynamics revealed by a coarse-grained modeling framework"

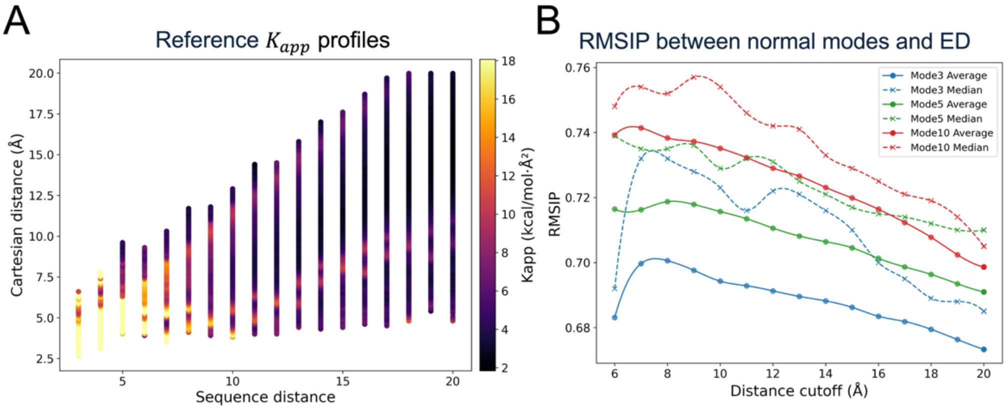


**Figure S1.** Parameter optimization of ICed-ENM. (*A*) Optimization of nonlinear spring model parameters from apparent force constant ($K_{app}$) profiles. Reference $K_{app}$ profiles were derived from the local dataset as functions of sequence distance ($d_{seq}$) and Cartesian distance ($d_{cart}$). The color scale represents the magnitude of $K_{app}$, reflecting effective inter-node stiffness inferred from time-ensemble fluctuations. These reference profiles served as training data for calibrating the nonlinear spring function in ICed-ENM. (*B*) Optimization of cutoff distance ($d_{cutoff}$) based on subspace similarity between ICed-ENM normal modes and MD-derived essential dynamics (ED). For the combined MD dataset (local and MDVerse datasets), the overlap between subspaces spanned by the lowest 3, 5, and 10 modes was quantified through the root-mean-square inner product (RMSIP), shown in blue, green, and red respectively. Mean and median RMSIP values are plotted as solid lines with circle markers and dashed lines with cross markers, respectively, as functions of $d_{cutoff}$.

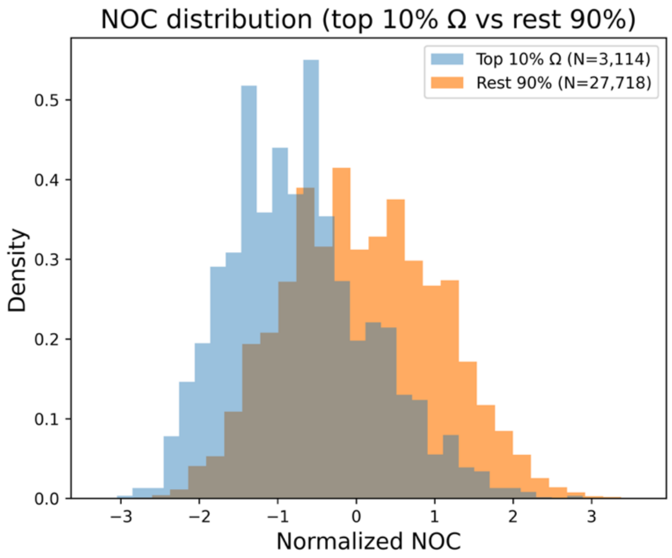


**Figure S2.** The distribution of residue-level number of contacts (NOC) for residues in the top 10% Ω group versus the remaining 90%. NOC was defined as the number of neighboring Cα atoms within an 8 Å cutoff from the target Cα and was Z-score-normalized per structure. The legend indicates the total number of residues in each Ω group.

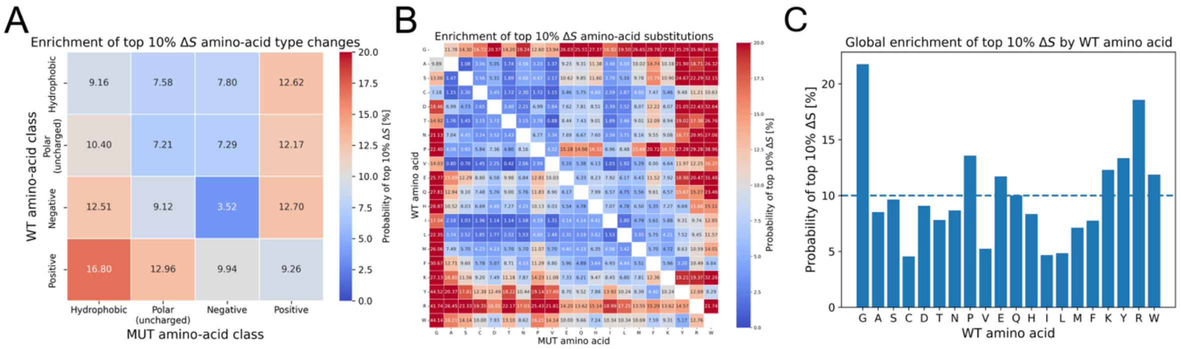


**Figure S3.** Amino-acid-type-dependent patterns of high $\Delta S$ mutations. The top $\Delta S$ group refers to mutations ranked within the top 10% of $\Delta S$ scores in each structure. (*A*) Probability of belonging to the top $\Delta S$ group as a function of the physicochemical classes of the WT and mutant amino acids (hydrophobic, polar uncharged, negative, and positive). (*B*) Probability of belonging to the top $\Delta S$ group across all 20 × 20 amino-acid pairs. For both heatmaps, the color scale is centered at 10%, corresponding to the background expectation. (*C*) Probability of belonging to the top $\Delta S$ group by WT amino-acid type. Bars represent the probability (%) that mutations originating from each WT residue belong to the top $\Delta S$ group. The dashed line marks the 10% background level.

**Table S1. Fundamental information about 5 proteins used in this study for unbiased MD simulations with different force-field models**

| Protein | PDB id[a], chain, status | Length[b] | Force-field model | Simulation time (*ns*)[c] |
|---|---|---|---|---|
| *Escherichia coli* Ribose-binding protein (RBP) | 1ba2, A, Open | 271 | AMBER99sb-ILDN (1), CHARMM36m (2), OPLS-AA/L (3) | 2 × 200 |
| | 2dri, A, Closed | 271 | AMBER99sb-ILDN, CHARMM36m, OPLS-AA/L | 2 × 200 |
| *Escherichia coli* 5'-nucleotidase (5'NTase) | 1hpu, A, Bound | 525 | AMBER99sb-ILDN, CHARMM36m, OPLS-AA/L | 2 × 200 |
| | 1oid, A, Unbound | 525 | AMBER99sb-ILDN, CHARMM36m, OPLS-AA/L | 2 × 200 |
| *Aquifex aeolicus* RNA endonuclease III (RNaseIII) | 1yyw, AB, Open | 438 | AMBER99sb-ILDN, CHARMM36m, OPLS-AA/L | 2 × 200 |
| | 1yyo, AB, Closed | 438 | AMBER99sb-ILDN, CHARMM36m, OPLS-AA/L | 2 × 200 |
| *Oryctolagus cuniculus* sarco(endo)plasmic calcium ATPase 1a pump (SERCA) | 2c9m, A, Open | 994 | AMBER99sb-ILDN, CHARMM36m, OPLS-AA/L | 2 × 200 |
| | 1t5s, A, Closed | 994 | AMBER99sb-ILDN, CHARMM36m, OPLS-AA/L | 2 × 200 |
| *Homo Sapiens* ATP-citrate lyase (ACLY) | 6pof*, ABCD, Open | 4384 | CHARMM36m | 3 × 200 |
| | 6hxh*, ABCD, Closed | 4384 | CHARMM36m | 3 × 200 |

[a] The asterisk indicates that the PDB model was re-modeled to add missing residues using SWISS-MODEL (4).
[b] Length indicates the number of residues of the pdb models.
[c] x × y indicates that simulation was performed for y ns with x replicas for each force-field model.

**Table S2. Dataset of unbiased MD simulations used in this study**

| Zenodo ID[a] | Protein | Length | Force-field model | Simulation time (*ns*) |
|---|---|---|---|---|
| 3991429 (5) | T4 lysozyme | 162 | AMBER ff15ipq with modified methyl rotation barriers (6) | 3 × 1 *μs* |
| 3989057 (7) | T4 lysozyme | 162 | AMBER ff99SB-ILDN with modified methyl rotation barriers (8) | 5 × 1 *μs* |
| 3754109 (9) | TonB protein from Helicobacter Pylori (30-285) | 256 | AMBER ff99SB-ILDN | 1 × 387 *ns* |
| 3778216 (10) | EN2 (143-259) | 117 | AMBER ff03ws (11) | 1 × 1 *μs* |
| 1010406 (12) | *Pseudomonas Aeruginosa* TonB-CTD (244-342) | 99 | AMBER ff99SB-ILDN | 1 × 400 *ns* |
| 3743358 (13) | *Helicobacter Pylori* HpTonB (30-285) | 256 | CHARMM36m | 1 × 400 *ns* |
| 6592231 (14) | GSH bound to Rat Multidrug resistance protein 1 (Mrp1) | 1339 | CHARMM36m | 1 × 1 *μs* |
| 6592231 (14) | GSSG bound to rat Mrp1 | 1339 | CHARMM36m | 1 × 1 *μs* |
| 6592231 (14) | Ritonavir bound to rat Mrp1 | 1339 | CHARMM36m | 1 × 1 *μs* |
| 6592231 (14) | GSH + Ritonavir bound to rat Mrp1 | 1339 | CHARMM36m | 1 × 1 *μs* |
| 6592231 (8) | GSSG + Ritonavir bound to rat Mrp1 | 1339 | CHARMM36m | 1 × 1 *μs* |
| 5226209 (15) | Glucagon-like peptide receptor (GLP-1R) + GLP-1 | 1163 | CHARMM36m | 4 × 500 *ns* |
| 5226209 (15) | GLP-1R + exendin-4 | 1164 | CHARMM36m | 4 × 500 *ns* |
| 5226209 (15) | GLP-1R + exendin-P5 | 1165 | CHARMM36m | 4 × 500 *ns* |
| 5226209 (15) | GLP-1R + oxyntomodulin | 1164 | CHARMM36m | 4 × 500 *ns* |
| 5811977 (16) | Meiosis 1-associated protein (M1AP) | 530 | CHARMM36m | 2 × 500 *ns* |
| 6558396 (17) | SARS-CoV-2 Spike RBD alpha variant (alpha-RBD) | 195 | CHARMM36m | 1 × 300 *ns* |
| 6558396 (17) | Alpha-RBD + BD23 | 423 | CHARMM36m | 1 × 300 *ns* |
| 6558396 (17) | Alpha-RBD + B38 | 629 | CHARMM36m | 1 × 300 *ns* |
| 6755131 (18) | Phenylalanine-4-hydroxylase (PAH) tetramer wildtype | 1808 | CHARMM36m | 1 × 100 *ns* |
| 6755131 (18) | PAH tetramer with muations | 1808 | CHARMM36m | 1 × 100 *ns* |

Those were retrieved from MDVerse data explorer (19).
[a] The trajectory data is accessible in the Zenodo repository: https://zenodo.org/records/[Zenodo ID].

**Table 3. Dataset of the transition PDB pairs**

| Open PDB id[a], chain, RG (Å) | Closed PDB id, chain, RG (Å) | Length | RMSD[b] (Å) | Collectivity[c] | Protein |
|---|---|---|---|---|---|
| 1ba2, A, 20.77 | 2dri, A, 19.08 | 271 | 6.28 | 0.77 | Ribose-binding protein (RBP) |
| 1hpuA, A, 24.32 | 1oid, A, 24.21 | 525 | 9.18 | 0.43 | 5'-nucleotidase (5'NTase) |
| 1yyw, AB, 26.27 | 1yyo, AB, 23.81 | 438 | 17.46 | 0.32 | RNA endonuclease III (RNaseIII) |
| 2c9m, A, 38.53 | 1t5s, A, 37.57 | 994 | 14.37 | 0.71 | Sarco(endo)plasmic calcuium ATPase 1a pump (SERCA) |
| 6pof*, ABCD, 62.98 | 6hxh*, ABCD, 59.75 | 4068* | 12.56 | 0.60 | ATP-citrate lyase (ACLY) |
| 2eia, B, 23.26 | 1eia, A, 22.88 | 204 | 8.07 | 0.67 | p26 capsid protein |
| 4ake, A, 19.46 | 1ake, A, 16.37 | 214 | 7.18 | 0.48 | *E.coli* Adenylate Kinase (ADK) |
| 1m8p, A, 30.15 | 1i2d, B, 28.13 | 572 | 4.74 | 0.63 | ATP Sulfurylase (ATPS) |
| 1dap, A, 22.00 | 3dap, B, 20.53 | 320 | 4.31 | 0.46 | Meso-diaminopimelic acid dehydrogenase (DAPDH) |
| 1dpe, A, 24.67 | 1dpp, A, 22.77 | 507 | 6.55 | 0.69 | Dipeptide binding protein (DppA) |
| 1ggg, A, 19.03 | 1wdn, A, 17.48 | 220 | 5.45 | 0.71 | Glutamine binding protein (GlnBP) |
| 2lao, A, 19.06 | 1lst, A, 17.70 | 238 | 4.74 | 0.69 | Lysine, arginine, ornithine (LAO) binding protein |
| 1lfg, A, 29.41 | 1lfh, A, 28.33 | 691 | 6.56 | 0.59 | Lactoferrin |
| 1rkm, A, 23.81 | 1jet, A, 22.90 | 517 | 3.16 | 0.64 | Oligo-peptide binding protein (OPPA) |
| 1tde, A, 21.34 | 1f6m, A, 21.24 | 315 | 7.35 | 0.67 | Thioredoxin reductase (TrxR) |
| 1fgu, A, 21.11 | 1jmc, A, 18.85 | 238 | 8.33 | 0.57 | ssDNA-binding domain of human replication protein A (RPA) |
| 1bp5, A, 21.26 | 1a8e, A, 19.28 | 328 | 6.75 | 0.77 | N-terminal lobe of transferrin |
| 1ram, B, 24.56 | 1lei, A, 23.11 | 273 | 3.11 | 0.65 | NF-kappa B p65 |
| 1ckm, A, 20.90 | 1ckm, B, 19.63 | 317 | 3.52 | 0.44 | mRNA capping enzyme |
| 1omp, A, 21.63 | 1anf, A, 20.72 | 369 | 3.78 | 0.73 | Maltodextrin-binding protein (mBP) |
| 1sx4, A, 29.23 | 1oel, A, 25.36 | 524 | 12.33 | 0.58 | Chaperonin GroEL |
| 1oao, D, 28.45 | 1oao, C, 27.03 | 728 | 7.07 | 0.66 | Acetyl-Coa synthase/carbon monoxide dehydrogenase |
| 6y2p, A, 27.32 | 7aex, A, 18.57 | 267 | 21.44 | 0.73 | mRNA endoribonuclease toxin LS |
| 7k6f, A, 25.65 | 7l6x, A, 20.33 | 250 | 14.48 | 0.70 | Transcription initiation factor TFIID subunit 1 |
| 6eze, B, 23.43 | 1ob2, A, 21.52 | 386 | 12.24 | 0.65 | *E.coli* Elongation factor Tu (EF-Tu) |
| 1tui, C, 23.54 | 1ob5, A, 21.60 | 397 | 11.67 | 0.65 | *T.aquaticus* Elongation factor Tu (EF-Tu) |
| 1kzq, A, 23.13 | 1ynt, F, 22.40 | 252 | 10.70 | 0.77 | Major surface antigen P30 |
| 7oh6, A, 38.58 | 7oh5, A, 37.68 | 1064 | 10.26 | 0.66 | Probable phospholipid-transporting ATPase DRS2 |

| | | | | | |
|---|---|---|---|---|---|
| 3njp, A, 34.86 | 1nql, A, 32.96 | 610 | 27.08 | 0.74 | Epidermal Growth Factor Receptor (EGFR) |
| 2rh5, A, 18.70 | 3sr0, B, 16.23 | 202 | 5.69 | 0.43 | *A.aeolicus* Adenylate Kinase (ADK) |
| 3k1p, A, 32.95 | 3k1p, B, 26.88 | 303 | 19.96 | 0.64 | *ADP1* HTH-type transcriptional regulator benM E266K mutant |
| 3tch, A, 24.33 | 3tcf, A, 23.14 | 517 | 5.16 | 0.67 | Periplasmic oligopeptide-binding protein (OppA) |
| 3b1o, B, 20.11 | 3b1o, A, 18.50 | 303 | 5.45 | 0.27 | Nucleoside kinase (NK) |
| 1jvk, A, 24.71 | 1jvk, B, 21.38 | 214 | 9.94 | 0.74 | Immunoglobulin lambda light chain |
| 4if4, A, 22.33 | 4gvp, C, 18.03 | 208 | 8.11 | 0.57 | Response regulator protein VraR |
| 4f48, A, 18.46 | 4f3h, A, 17.68 | 241 | 4.86 | 0.07 | FimX protein |
| 1iqp, C, 24.58 | 1iqp, A, 24.04 | 319 | 8.21 | 0.24 | Replication fator-C small subunit (RFCS) |
| 2xro, B, 22.10 | 2xrn, A, 19.57 | 239 | 12.33 | 0.64 | Multidrug-binding protein TtgV |
| 3fds, A, 25.80 | 1n48, A, 22.38 | 341 | 15.38 | 0.40 | DNA polymerase IV |
| 2h6c, B, 27.25 | 2h6c, A, 24.15 | 208 | 9.68 | 0.59 | Oxidized chlorophenol reduction gene K (CprK) |
| 1iz1, B, 32.28 | 1iz1, A, 23.53 | 292 | 18.11 | 0.72 | CbnR |

[a] The asterisk indicates that the PDB model has missing residues.

[b] RMSD quantifies the spatial difference between two given PDB structures.

[c] The value of collectivity quantifies how globally the structural changes occur.

16. U. Gerlevik, M. C. Ergoren, O. U. Sezerman, S. G. Temel, All-atom Molecular Dynamics

Simulations of Meiosis 1-associated protein (M1AP) to Investagate the Impact of Known Missense Mutations Associated with Male Infertility through Non-obstructive Azoospermia. Zenodo. https://doi.org/10.5281/ZENODO.5811977. Deposited 1 March 2021.

17. D. Ray, R. N. Quijano, I. Andricioaei, Simulation of Receptor Binding Domain of SARS-CoV-2 spike protein (WT and variants) in complex with neutralizing antibodies. Zenodo. https://doi.org/10.5281/ZENODO.6558396. Deposited 17 May 2022.

18. T. Aslan, *et al.*, All-atom molecular dynamics simulations of phenylalanine-4-hydroxylase (PAH) tetramer to investigate the impact of two novel heterozygous mutations, p.Y198N and p.Y204F, observed in a classical phenylketonuria patient. Zenodo. https://doi.org/10.5281/ZENODO.6755131. Deposited 26 June 2022.

19. J. K. Tiemann, *et al.*, MDverse, shedding light on the dark matter of molecular dynamics simulations. *eLife* **12**, RP90061 (2024).
